# Microbial map of the world’s vineyards: Applying the concept of microbial *terroir* on a global scale

**DOI:** 10.1101/2020.09.25.313288

**Authors:** A. Gobbi, A. Acedo, N. Imam, R.G. Santini, R. Ortiz-Álvarez, L. Ellegaard-Jensen, I. Belda, L.H. Hansen

## Abstract

The specific microbial biodiversity linked to a particular vineyard location is reported to be a crucial aspect, in conjunction with edaphic, climatic and human factors, in the concept of wine *terroir*. These biogeographical patterns are known as microbial *terroirs*.

This study applied an HTS amplicon library approach in order to conduct a global survey of vineyards’ soil microbial communities. In all, soil samples from 200 vineyards on four continents were analysed in an attempt to establish the basis for the development of a vineyard soil microbiome map to represent microbial wine *terroirs* on a global scale.

This study established links between vineyard locations and microbial biodiversity on different scales: between continents and countries, and between different wine regions within the same country. Geography had a strong effect on the composition of microbial communities on a global scale, which was also maintained on a country scale. Furthermore, a predictive model was developed, based on random forest analyses, to discriminate between microbial patterns in order to identify the geographical source of the samples with reasonable precision. Finally this study is the first to describe the microbial community of new and northern wine-producing regions, such as Denmark, that could be of great interest for viticulture adaptation in a context of climate change.

## Introduction

Wine is a multi-billion dollar market of high cultural and economic value (Bokulich et al. 2016). Since the start of wine history (Zohary 2003), place of origin is a major factor driving wine purchase decisions, followed by competitive prices and brands. However, this by itself is not always sufficient to judge a wine’s identity or value (Ballantyne et al. 2019). Winemakers rely on the concept of *terroir* to explain the unicity of their wine in terms of taste and flavour. This term was originally used in Burgundy in the 1930s as a marketing tool to differentiate between wines (Whalen 2009), but it now goes beyond the wine sector and is used to explain the distinctive regional characteristics of other high-value products, especially those where microbial fermentation has taken place, such as cheese, coffee and cocoa (Østerlie and Wicklund 2018; Planète Terroirs 2010)

Today, the concept of wine *terroir* has spread around the world, and wine-producing countries are trying to regulate it with the legal definition of appellations of origin. For example, 139 American viticultural areas have been recognised in California (USA) alone (Wine Institute, 2020) and 90 in Spain (Agricultura y Ambiente MAPA 2016). In this context, it is well known that certain locations have optimal climate and soil characteristics for the production of premium wines, thus the appellation of origin system plays an important role in the legal classification of wines, as well as in the final customers’ purchase decision. Consequently, protecting the integrity of this classification system is of paramount importance to producers, distributors, retailers and, of course, consumers (Ballantyne et al. 2019). Thus, a major aim – and an exciting research area – is to establish the scientific basis of wine *terroir*, which relies on a multitude of dimensions such as local edaphic, climatic, human and biotic factors that contribute to modifying the quality and the traits of the resulting wines (OIV/VITI 333/2010). Among them, the specific microbial biodiversity associated with a certain vineyard location is reported to be a key aspect associated with biogeographical patterns and directly involved in vine growing, grape quality and winemaking (Belda et al. 2017).

In the wine *milieu*, pioneering studies based on high-throughput sequencing (HTS) have been conducted by Bokulich et al. (2012), Burns et al. (2016) and Zarraonaindia et al. (2015) that have revealed microbial biogeography associations across multiple viticulture areas. These biogeographical patterns have subsequently been confirmed in other places such as Catalonia in Spain (Portillo et al. 2016) and in Italy’s Cannonau wine region (Mezzasalma et al. 2017). Microbial biodiversity also provides relevant insights into the impacts of agricultural management and soil quality (OIV 2018; Hermans et al. 2020).

Furthermore, Bokulich et al. (2016) and Knight et al. (2015) have also shed light on the associations between the microbial and the metabolic fingerprint of wine produced in different wine regions. In this field, Belda et al. (2016) has also described distinctive and clustered metabolic patterns for wine-related yeasts depending on their geographical origin. The links between the unicity of a wine’s metabolic profile and all the factors affecting the grapes are known and valuable to the viticulturist, supporting the ancient concept of *terroir*. Within this complex and multifaceted concept, the microbiome of vineyards has been shown to be a unique and integrative biomarker (Belda et al. 2017; Gilbert, van der Lelie and Zarraonaindia 2014) that affects wine quality indirectly (by affecting vine health and physiology), but also directly as the main reservoir of autochthonous fermentative microbiota. A comprehensive review on this topic is given in a book chapter by Belda et al. (2020).

The role of microbiota in all stages of the wine production process, the presence of soil as the main reservoir for microbes and the non-random microbial-geographical association reveal the potential applied impact of microbial *terroirs*. This could serve as a biological objective for future biotechnological applications regarding productivity or disease resistance (van der Heijden and Hartmann 2016), but also contribute to defining biomarkers for monitoring and protecting the biological determinants of wine regions. In this context, soil biodiversity remains one of the most recognised parameters linked to the concept of sustainable agriculture (Altieri 1999; Brussaard, de Ruiter and Brown 2007; Nielsen, Wall and Six 2015). Therefore understanding global patterns in the microbial community composition of vineyard soils worldwide provides the information required for future advances in the sustainability of viti-viniculture, since wine grapes are one of the most dramatically affected crops in the current global change scenario (OIV Expertise 2018).

This study applied an HTS amplicon library approach to conduct a global survey of the soil microbial communities of vineyards in 13 countries, including different wine regions and weather conditions, in an attempt to establish the basis for the development of vineyard soil microbiome maps that, with further modelling efforts on different scales, will provide a better understanding of the role of microbes in connecting vineyard *terroir* with wine quality. The aim of this study was to demonstrate that the link between distinctive microbial communities and specific wine regions is a concept that can be extended to a global scale, allowing a better understanding of the biogeographical basis and the microbiome boundaries of vitivinicultural *terroirs*. Finally, for a statistical demonstration of the microbiome fingerprint within different wine regions worldwide, a random forest-based predictive model was built in order to trace the origin of a given soil sample based on its microbial community composition.

## 2 Materials and Methods

### 2.1 Materials

This study involved a microbial amplicon-based survey created with previously unpublished data combined from two different datasets. The data in this study originated partly from the MICROWINE project but also included private data from Biome Makers Inc. obtained with BeCrop® Technology. This project involved a total of 252 soil samples from 200 vineyards. These samples were therefore collected by different people around the globe between 2015 and 2018. Although there were some differences in the sampling scheme, storage conditions and sample processing, a general description of this protocol can be outlined as follows. All the samples were of topsoil taken at a depth of between 0 cm and 10 cm. The MICROWINE samples consisted of five samples collected and sequenced for each field, while the Biome Makers samples pooled together topsoil from three random spots in each field with the DNA then extracted from this composite sample. The shipment conditions of the soil samples were either −20 °C or non-refrigerated, followed by −20 °C or −80 °C as the storage temperature in the laboratories until DNA extraction. DNA extractions were performed using bead beating-based DNA extraction kits such as the Dneasy Powerlyzer Powersoil Kit (Qiagen) for the BeCrop® platform (patent publication number: WO2017096385, Biome Makers) and the FAST-DNA Spin Kit for Soil (MP-Biochemical) for the MICROWINE project. A complete overview of all the samples used in this study and their relative origin is given in Table S1. The use of bead beating-based kits, such as those included in this study, ensured that the results produced were comparable and also allowed the recovery of the highest biodiversity within soil samples (Vishnivetskaya et al. 2014).

### 2.2 Dataset

This study analysed soil samples from 200 locations (in 13 countries on four different continents) that had all been collected close to the harvest period. This allowed a dataset to be built that was strongly dependent on the geography and included vintage-related parameters such as average maximum and minimum temperatures and precipitation measured for the two weeks prior to sample collection. Analyses of the microbial composition on a country-level scale was performed using the samples from Spain, the country with the highest number of samples (n=86) from 12 wine regions.

Weather information was retrieved from World Weather Archive (https://www.worldweatheronline.com/) with the closest reference point chosen based on the GIS coordinates identifying the samples.

The data are publicly available on ENA under the following study accession number: PRJEB40350

### 2.3 Library preparation

The general protocol for the library preparation is outlined below, although some modifications occurred between studies. All the libraries were prepared following the two-step PCR Illumina protocol and these were subsequently sequenced on an Illumina MiSeq instrument (Illumina, San Diego, CA, USA) using 2×251 paired-end reads for the MICROWINE samples and 2×301 paired-end reads for the Biome Makers samples.

The same variable regions were investigated in both datasets (MICROWINE and Biome-Makers) for 16S and ITS with a few differences in the primer sequences. A complete description of the primers is given in Table S2. All the PCR reactions were prepared using UV-sterilised equipment and negative controls were run alongside the samples. Furthermore, PCR conditions, such as the number of cycles, annealing temperature, thermocycler and Master Mix composition, changed between samples from different projects. The library for both datasets was prepared using a two-step PCR, as described by. Gobbi et al. (2019) with modifications. In the MICROWINE dataset, the sample concentration was approximately 10-20 ng of DNA, and both PCRs were carried out using a Veriti Thermal Cycler (Applied Biosystems). In each reaction in the first PCR for 16S, the mix contained 12 μL of AccuPrime™ SuperMix II (Thermo Scientific™), 0.5 μL each of forward and reverse primer from a 10 μM stock, 0.5 μL of bovine serum albumin (BSA) to a final concentration of 0.025 mg/mL, 1.5 μL of sterile water and 5 μL of template. The reaction mixture was pre-incubated at 95 °C for 2 min, followed by 33 (16S) and 40 (ITS) cycles of 95 °C for 15 sec, 55 °C for 15 sec, 68 °C for 40 sec, and then a final extension at 68 °C for 4 min. The samples were subsequently indexed by a second PCR using the protocol in Feld et al. (2015). Finally, the samples were pooled in an equimolar amount before sequencing. Biome Makers samples were obtained amplifying the 16s rRNA V4 region, while the ITS were obtained by amplifying the ITS1 region using BeCrop® custom primers (patent WO2017096385).

### 2.4 Bioinformatics

All the data produced and collected were subsequently analysed using QIIME2 v2019.7 (Bolyen et al. 2018), as described in Gobbi et al. (2020). Reads from the 16S rRNA gene produced following the MICROWINE and Biome Makers protocol were collected and processed using DADA2 single read analyses (Callahan, McMurdie and Holmes 2017). The phylogenetic tree was calculated based on the insertion fragment plugin to reduce the batch effect from using two different primer sets (Janssen et al. 2018). Taxonomy was assigned using a Naive Bayes classifier (Bokulich et al. 2018) trained with Greengenes v13_8 (DeSantis et al. 2006).

ITS sequences were analysed using a DADA2 single read without merging. Denoised reads were used to build a phylogenetic tree using MAFFT (Katoh and Standley 2013) and subsequently taxonomy was assigned using a Naive Bayes classifier trained with the UNITE database (full alignment) (Nilsson et al. 2019).

The frequency tables for 16S and ITS were rarefied to 10000 high-quality reads per table and used for all subsequent analyses of diversity and composition.

### 2.5 Statistics

Alpha diversity was calculated using the Shannon Index and tested with a Kruskal-Wallis test with adjustment for multiple testing. Beta diversity was calculated based on unweighted UNIFRAC (Lozupone and Knight 2005). Kruskal’s non-metric scaling was used to perform a principal coordinate analysis based on UNIFRAC distances between samples. The results were plotted, labelling samples by country and continent. These groupings were tested with PERMANOVA. To assess the global core microbiome, taxa were pre-selected if they were detected in each continent. This was visualised as a Venn diagram using the VennDiagram R package (Chen 2018). These “core” taxa were evaluated on a continuum of abundances and prevalences, and plotted as a heatmap as per Salonen et al. (2012).

For subsequent analyses, taxa that had fewer than 20 non-zero abundances in the whole dataset were aggregated into an “others” variable. A zero replacement was also performed using Bayesian inference with a Dirichlet prior and multiplicative adjustment to maintain other proportions (Martín-Fernández et al. 2015).

A redundancy analysis (RDA) was performed using a centred log ratio (CLR) of 16S and ITS abundances, constrained by the country or continent from where the samples came. Additionally, weather conditions (specifically average minimum and maximum temperatures and precipitation) at sampling were used as conditioning variables for RDA. The RDA analysis was conducted using the vegan R package (Oksanen et al. 2019).

The predictive potential of the microbiome for geographical region was assessed using random forests. Models were fitted using CLR-transformed 16S and ITS abundances as predictors and country or continent as outcome. Additional random forests were fitted using weather variables (as mentioned above) in conjunction with abundances as predictors. Subsequently, 75 % of the dataset was randomly selected and used for repeated cross-validation, using a grid of values for hyper-parameters (Supplementary Table S6). Models were selected for the highest accuracy in the test set (remaining 25 % of the dataset). Country and continent-level confusion matrices were generated for the test set (Table S4). Feature ranking was performed using the Gini index. Random forests were fitted using the ranger R package (Wright and Ziegler 2017). All the analyses were performed in the R programming environment (R Core Team 2019) and Qiime2 v2019.7 (Bolyen et al. 2018).

## 3 Results

### 3.1 Study coverage and taxonomic features of the global vineyard soil microbiome

This work provided new insights into the microbial community of vineyard soils worldwide by analysing amplicon data for bacterial 16S rRNA gene and fungal ITS region sequencing. The resulting database was explored from a multi-scale perspective to understand the impact of geography and weather on the microbial community composition in different wine-producing countries.

The sequencing dataset obtained was representative for 13 different countries on four continents with the intention of covering most of the dominant areas for grapevine cultivation (Fig. 1). These 13 countries represent more than 83 % of total wine production worldwide (OIV 2019). In all, 252 soil samples from 200 different vineyard blocks were sampled close to harvest time, resulting in a total of 504 analysed samples equally divided between 16S and ITS amplicon sequencing.

**Figure 1.**
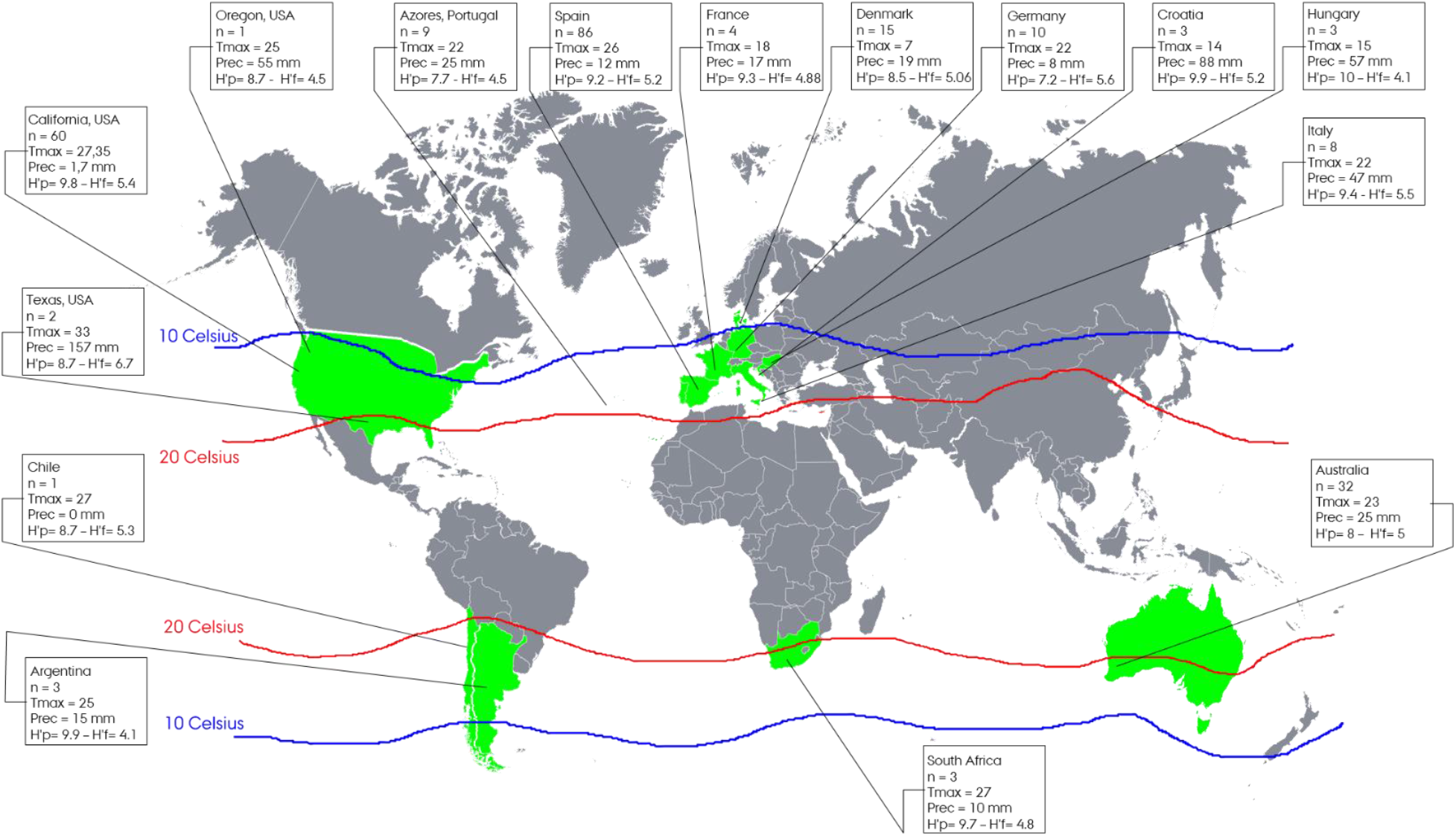
Coverage of the study. The countries highlighted in green are represented in the study’s dataset. The two isothermal lines define the range for optimal conditions for grapevine cultivation (isothermal source: https://www.thirtyfifty.co.uk/spotlight-climate-change.asp). Each panel shows the country, the number of samples and information about the average maximum temperature and the level of precipitation up to two weeks before sample collection. Finally, H’P and H’F define the Shannon Index for the prokaryotic and fungal community respectively.

After analysing the data, 84770 ESV for 16S and 33254 for ITS were obtained. Based on their taxonomical annotations, among the 45 prokaryotic phyla identified in these samples, 12 showed relative abundances above 1 % in at least one of the 13 countries. *Proteobacteria* occurred with the highest relative abundances in all 13 countries, with values varying between 18.8 % (Portugal) and 32.5 % (Argentina). *Actinobacteria* were the second highest abundant bacterial phylum, ranging from 5.9 % (Croatia) to 28.4 % (Portugal) in most of the countries except Croatia, Germany and Italy where *Acidobacteria* replaced *Actinobacteria* as the second highest abundant bacterial phylum, while elsewhere it was the third most represented phylum. Additionally, *Planctomycetes* (Spain and USA)*, Bacteroidetes* (France, Hungary and Croatia) and *Chloroflexi* (Portugal) showed a relative abundance above 10 %, and together with *Verrucomicrobia, Firmicutes* and *Gemmatimonadetes* consistently showed a considerable relative abundance, with values above 5 %. The archaea phylum *Crenarchaeota* (dominant genus *Nitrososphaera*) showed a large relative abundance variation in the 13 countries, with the main values detected in Portugal (10.9 %), Chile (12.5 %) and Germany (29.5 %), while the values in the other countries ranged between 1.6 % and 8.2 %.

In the evaluation of fungal taxa detected with relative abundances above 1 %, *Solicoccozyma (sin. Cryptococcus)*, when present, was the dominant genus in eight of the 13 countries. The relative abundance of *Solicoccozyma* in Argentina, Chile, South Africa, Italy and Croatia ranged from 13.4 % to 39.3 %. In the countries where *Solicoccozyma* was not dominant (Australia, Denmark, Germany and Portugal), *Fusarium* and *Cladosporium* were the most dominant, except in the USA. In Portugal and South Africa, *Fusarium* had a relative abundance up to 10 %. *Mortierella* was identified in all countries, but in six of them the relative abundance values were above 5 %. Other abundant fungal taxa were *Alternaria* in Chile (17.6 %) and South Africa (9.2 %), *Aspergillus* (often above 1 % and up to 5 %), and *Gibberella intricans* (Portugal, Denmark and Spain, where it reached up to 9.3 %). Some taxa appeared to be dominant in just one or two countries, for instance *Metarhizium_robertsii* (9.14 % in Denmark), a *Basidiomycota-*like *Lepiota farinolens* in South Africa (13.6 %), *Sclerotinia sclerotiorum* (12.8 %) and *Rhizopus arrhizus* (13.9 %) in France, *Tausonia pullulans* in Hungary (7.5 %) and Spain (2.2 %), and *Truncatella* in Germany (6 %). *Xylariales* in Portugal (17.8 %) was assigned to *Robillarda sessilis*, although it may represent a different species since after blasting the read the identity was only 98 %; the same was found for *Idriella* in Argentina (16 %), *Chaetothyriales* (6.3 %) and *Rozellomycota* (7.8 %) in Croatia and *Onygenales* (8.8%) in France. All these results are given in Figure S1a and Figure S1b.

Since the sample count was not evenly distributed between the different countries, alpha diversity was measured using the Shannon Index because it is more tolerant to the sampling coverage given its statistical nature. From this analysis, displayed in Figure 1, the Shannon Index for prokaryotes (H’P) and fungi (H’F) are presented here as the average value for each country. H’P ranged from 7.2 (Germany) to 9.9 (Hungary, Croatia and Argentina), while H’F ranged from 4.1 (Hungary and Argentina) to 6.7 (Texas, USA) and was therefore consistently in lower ranges than H’P in all the different countries. An analysis was performed on possible correlations between the Shannon Index in all the different samples and some meteorological parameters such as the average maximum temperature and precipitation data prior to sample collection. No strong relationship was identified with these parameters, suggesting that geography or other edaphic factors are more dominant features than weather conditions, which depend more on latitude than longitude.

### 3.2 Structure of microbial communities in vineyard soils: beta diversity and core microbiota

The results outlined in this section identified how similar and how diverse the communities were between different areas, as summarised in Figure 2 and Figure 3. The beta diversity was analysed and results visualised using a PCoA plot based on CLR on global and national scales. Based on the current visualisation, there was a consistent clusterisation for the individual countries represented in the dataset in both the prokaryotic (Fig. 2a) and fungal communities (Fig. 2b). At national scale all the samples came from Spain, the most represented country (n=86). The cluster distribution once again highlighted the different regions sampled (Fig. 2c and 2d).

**Figure 2.**
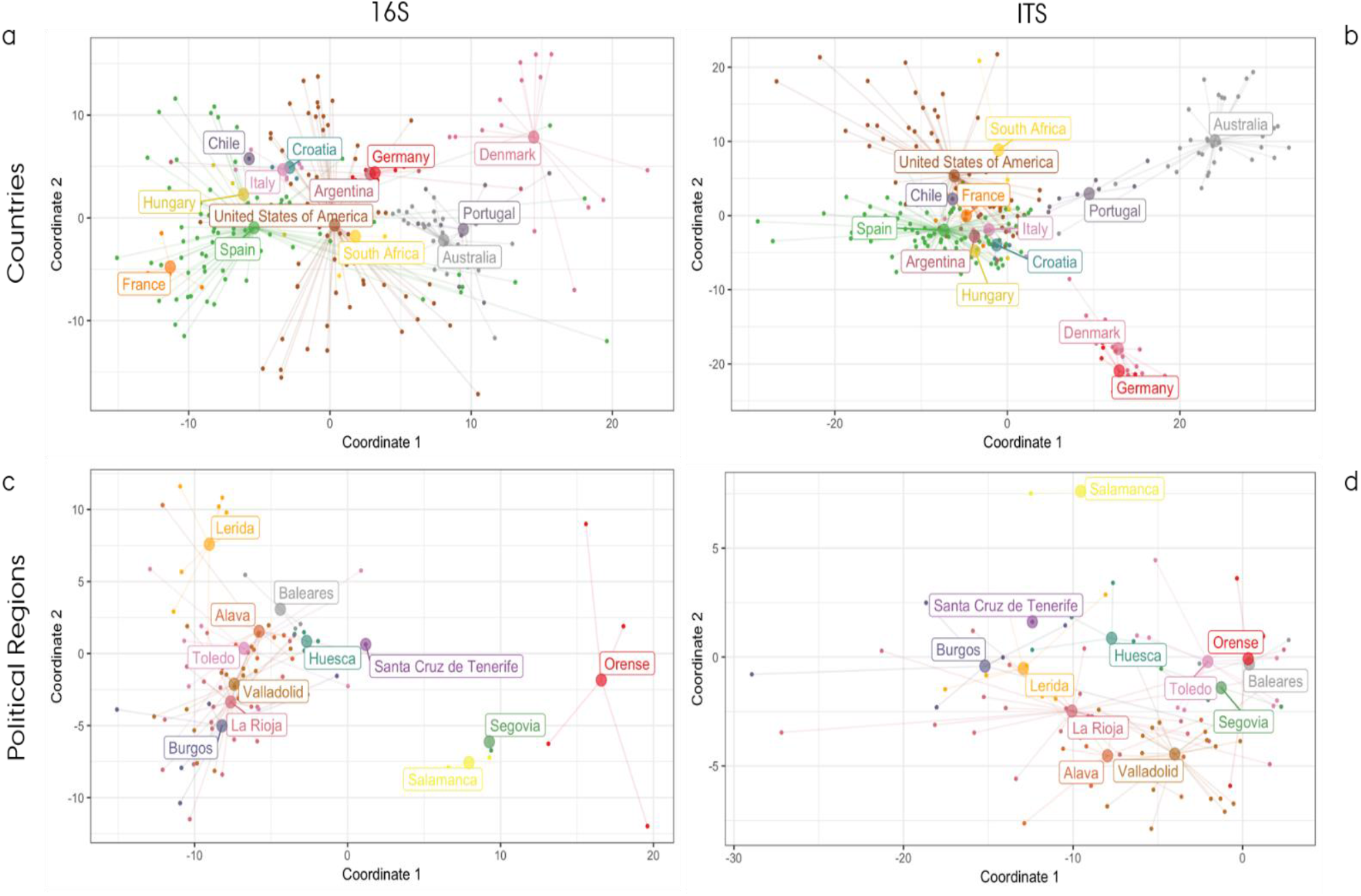
PCoA plots of beta diversity based on CLR. Figures 2a and 2b represent the national-scale distribution of prokaryotes (2a) and the fungal community (2b); Figures 2c and 2d represent a regional-scale distribution for prokaryotes (2c) and fungi (2d) on a subset of samples from Spain.

**Figure 3.**
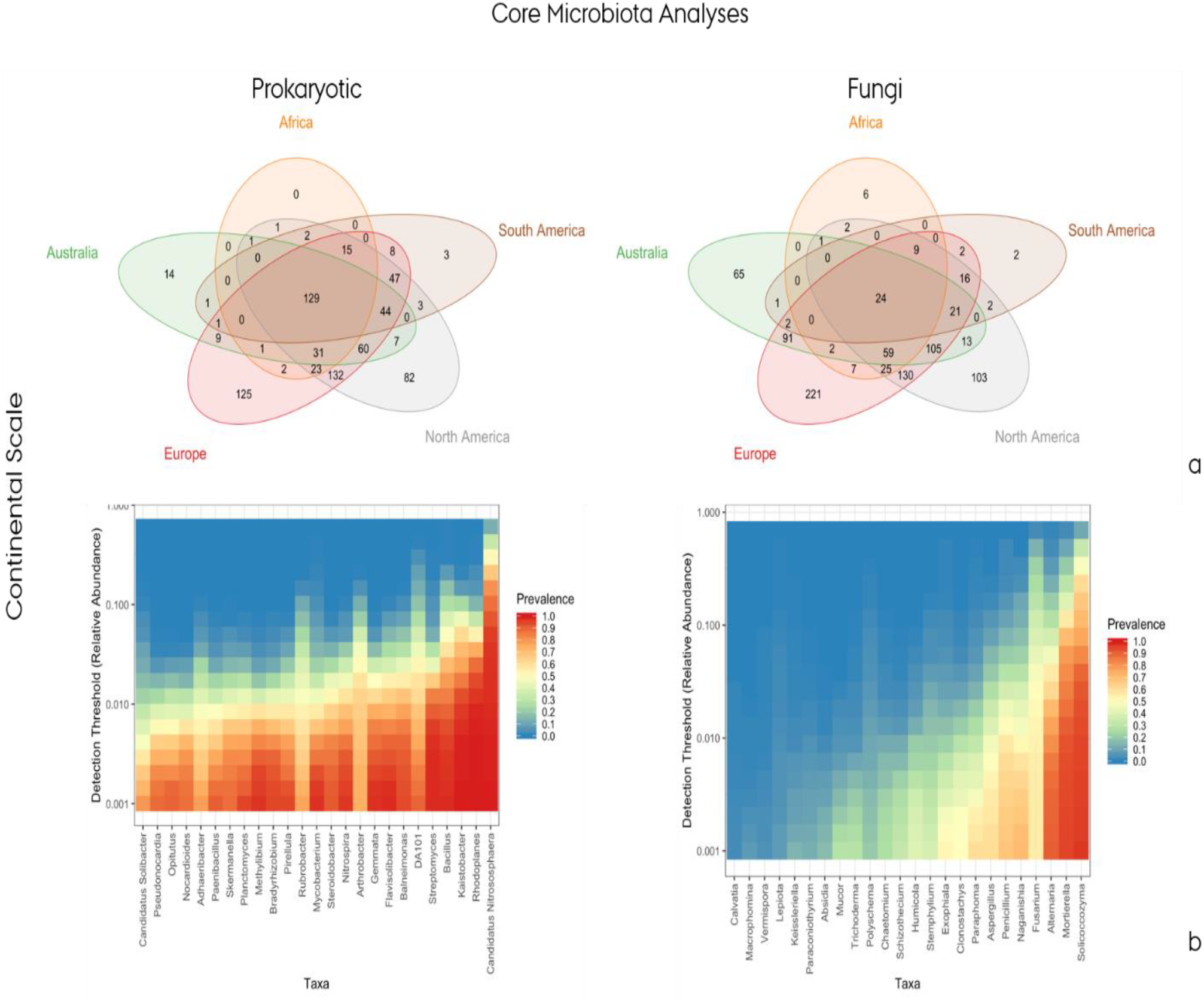
Evaluation of core microbiota: a) co-occurrence of different genera at continent level for the prokaryotic (left) and fungal (right) community; b) heatmaps for the top 25 genera shared between continents for the prokaryotic (left) and fungal communities (right), including information on their relative abundance.

The effect of geography on different scales was measured using PERMANOVA. The results were consistent with the initial hypothesis about the impact of geography on microbial communities in vineyard soils. This was more evident when reducing the scale from a global comparison to both a continent (prokaryotic: R2=0.08, p=0.001; fungal: R2=0.09, p=0.001) and country level (prokaryotic: R2=0.16, p=0.001; fungal: R2=0.21, p=0.001), and then to a national-scale comparison between regions (prokaryotic: R2=0.27, p=0.001; fungal: R2=0.25, p=0.001) (Table S3).

Although the impact of geography on a large scale was important overall, it was not consistently significant in a few neighbouring underrepresented countries (i.e. Italy and Croatia for 16S or Germany and Denmark for ITS). This could be due to the relative high variance between the microbial community retrieved from different vineyards in the same country. Therefore, in order to improve the resolution of the analyses, meteorological data were included as an additional constraint. This effect could be seen when looking at the variance partitioning RDA analyses performed (displayed in Fig. S2), where the use of an additional constraint helped resolve closely related countries and contemporarily reduced the batch effect. However, as observed in the abovementioned PERMANOVA analysis, it is important to highlight that the variance explained by weather data (average maximum and minimum temperatures and precipitation) in the composition of microbial communities was lower than the variance explained by geography, irrespective of the scale (Table S3). In this context, it can be assumed that this approach would given an even higher resolving power if additional constraints (e.g. soil properties) were considered.

The evaluation of the core microbiota, which were stable in the soil in these vineyards regardless of their geographical distance, produced a list of ubiquitous genera (129 prokaryotic and 24 fungal) present in 100 % of the countries and in at least 80 % of all the samples from each country (Fig. 3a). From this list of core taxa, the global prevalence was explored, at different relative abundance levels, of the most widespread distributed genera among the prokaryotic and fungal populations (Fig. 3b). Among the dominant prokaryotic genera were *Nitrososphaera* (Archaea), *Rhodoplanes, Kaistobacter, Bacillus* and *Streptomyces* due to their large prevalence at relative abundance values higher than 1 %. Similarly, the main dominant representatives of the core fungal communities were *Solicoccozyma, Morteriella* and *Alternaria*. Figure 3b suggests that there was a more diverse and balanced core within the dominant bacteria genera than within the fungal core microbiota, where just three genera seemed to dominate the populations in a substantial proportion of the samples.

### 3.3 Random forest

For a practical demonstration of the effect of geography in determining the structure of microbial communities in vineyard soils as a key consideration when defining wine *terroirs*, a predictive model was developed based on random forest analyses. The objective was to trace the origin of a certain soil sample based solely on its microbiome composition. The fitted models for each level (country or continent), sequencing type (ITS or 16S) and inclusion of weather variables had test set accuracies of between 80.0 % and 93.3 %. Test set accuracies for the final models are presented in Table S5. Intuitively, the continent level models were less accurate regardless of the inclusion of weather variables included in the analysis. However, quite notably, the inclusion of weather variables did not seem to have a large effect on test set accuracy when used in conjunction with sequencing data.

Figure 4 shows the confusion matrix reporting the predictions from the trained models. As mentioned above, the reasonable accuracy of the fitted models resulted in a high rate of coincidence between the actual and predicted origins of the samples, with a similar performance of prokaryotic and fungal-based models. Figure 4b shows the taxonomical identity of the 20 best predictor phylotypes. As mentioned above, weather data did not have a large effect on test set accuracy, and microbial variables remained the main predictors in these mixed models (Fig. S3).

**Figure 4.**
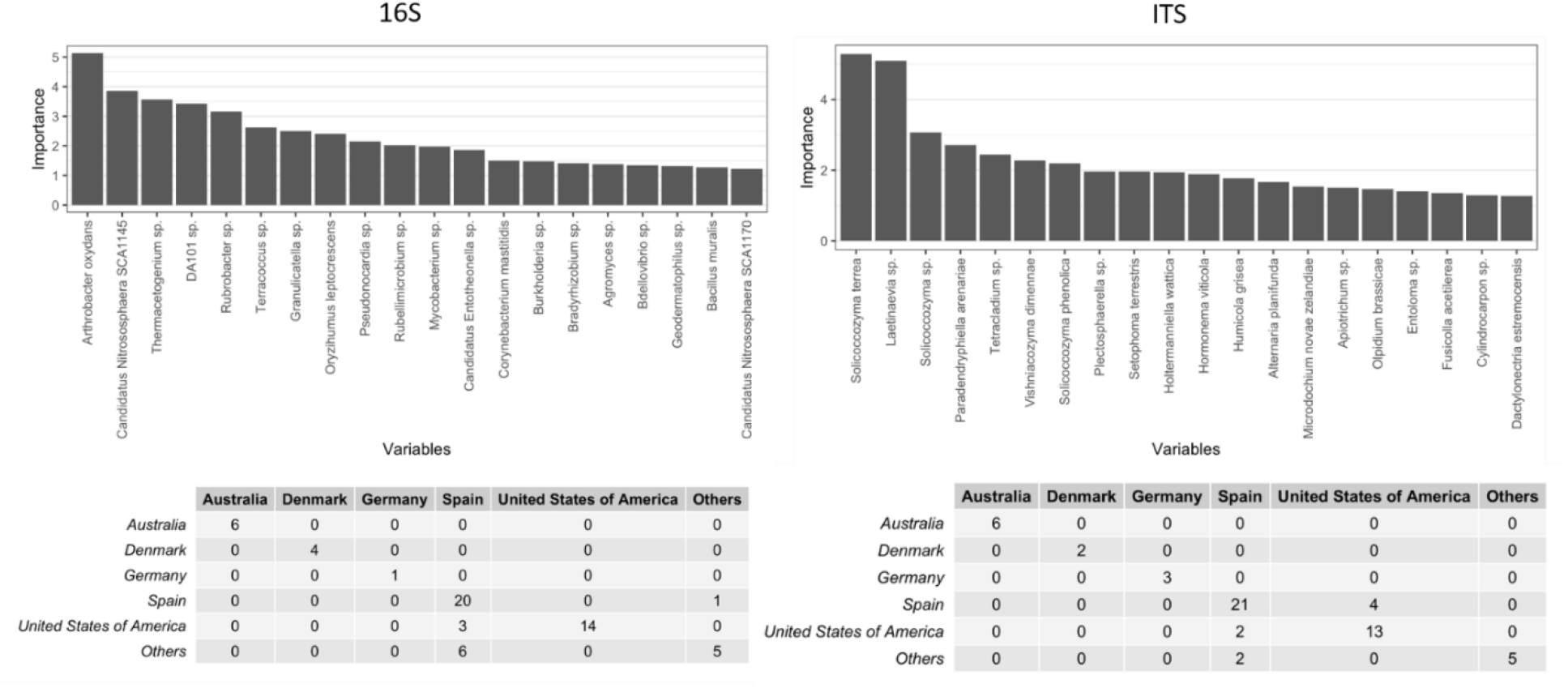
Random forest results at national scale including only microbial data. The top charts shows the confusion matrixes for the prokaryotic and fungal communities; the bottom charts give the 20 best predictor for prokaryotic (right) and fungal (left) community.

## 4 Discussion

### 4.1 Database and batch effect

When combining datasets from different studies, the impact of what is known as batch-effect bias needs to be addressed. The database in this study was composed of samples from several countries (see the detailed list of samples included in this study in Table S1), and although they were produced in a similar way, confounding factors could be expected in the global distribution. Batch effect is a very common bias, especially in meta-analytical studies (Henschel, Anwar and Manohar 2015), that can influence the results and subsequently the conclusions. This risk is even higher when the same type of sample is compared, while the effect is lower when different types of samples are analysed together, as reported by several meta-analyses (Henschel, Anwar and Manohar 2015; Lozupone et al. 2013; Lozupone and Knight 2007). With regard to the 16s rRNA gene analyses, an algorithm called SEPP (S. Janssen et al. 2018) was chosen that has been shown to dramatically reduce the bias due to the use of different primers or to the amplification of different hypervariable regions. This method is not applicable to ITS since the lack of a high-quality phylogenetic tree will affect the results more than a traditional pipeline. For ITS, a single-read DADA2 approach with basic filters was used instead that has been shown to represent one of the best approaches for amplicons so far (Pauvert et al. 2019) and has also previously been used (Alex Gobbi et al. 2020).

Thanks to this analytical effort, and even assuming a certain bias, the effect of geography in the composition and structure of microbial communities was still visible (Fig. 2) and statistically relevant (p-value < 0.001). On a global scale, alpha diversity was measured using the Shannon Index, which is relatively stable against the batch effect from the different primers used. This is because it also accounts for evenness in the ESV distribution and is not solely based on their presence/absence (Engelbrektson et al. 2010). Furthermore, once singletons are removed, its entropy coverage adjustment accounts for unobserved taxa caused by an uneven coverage of the countries (Willis 2019). Another approach is to compare the co-occurrence pattern within the whole database after taxonomy assignment at different taxonomical ranks (Engelbrektson et al. 2010). In fact, two EVS that have a different nucleotide sequence, coming from different regions of the same gene marker, could be assigned to the same taxa, despite some differences that can occur in relative abundance.

### 4.2 Microbial diversity in global vineyard soils

The PCoA plots confirmed the link between geography and the microbial community in vineyards on global and regional scales. There has been some limited evidence of this on a regional scale, as already demonstrated by Burns et al. (2015) and Bokulich et al. (2014) in California, Knight et al. (2015) in New Zealand, Castañeda and Barbosa (2017) in Chile, and Coller et al. (2019) and Mezzasalma et al. (2017) in Italy. The present study also demonstrated that the concept of microbial *terroir*, previously established by Gilbert, van der Lelie and Zarraonaindia (2014), can be scaled up to a global scenario, since geography has a strong effect on shaping the microbial community of vineyard soils from different countries, with just a few clusters overlapping. This led to the suggestion that there is a hierarchy of geographical distances, with the general trend being the further the distance, the more diverse the community. The present study confirmed the regional-scale clusterisation when looking at the samples from Spain (Fig. 2c and 2d).

This was the case for bacteria and fungi on an international scale, but was even more noteworthy when looking at the distribution within the same country. This suggests that the use of microbial information is a sensitive parameter for discriminating between more closely located communities when confounding factors are reduced. In contrast, the performance of the random forest model displayed an opposite trend, with greater accuracy on a continental scale than on a national scale. Moreover, the accuracy seemed to be greater when looking at mycobiota. This is possibly due to the fact that the ITS gene marker may give a deeper resolution than the bacterial 16S rRNA gene marker. However, the study of bacterial populations is more widely referenced nowadays than the fungal community, which suffers from a lack of curated and updated databases or an optimised algorithm and pipeline to infer phylogenetic relations between fragments of variable length (Pauvert et al. 2019).

At a taxonomic level, *Proteobacteria* and *Acidobacteria* have previously been identified as the most abundant bacterial phyla within soils (P. H. Janssen 2006), including topsoil studies carried out in vineyards (Burns et al. 2015; Zarraonaindia et al. 2015). In agreement with such observations, *Proteobacteria* was the most abundant phylum in all 13 countries evaluated; however, instead of *Acidobacteria* other phyla such as *Actinobacteria* occurred in higher relative abundances.

The widespread occurrence of *Crenarchaeota* members in these samples is in agreement with the detection of this archaeal phylum in a range of environments around the world, such as agricultural fields, sandy soil, forest soil, contaminated soil and the rhizosphere (Bintrim et al. 1997; Simon, Dodsworth, and Goodman 2000). However, while the relative abundance fraction of this phylum is typically reported as representing up to 5 % of the total prokaryotic community (Buckley, Graber and Schmidt 1998; Ochsenreiter et al. 2003), it was found in 10 of the 13 countries in values exceeding 5 %, with an emphasis on samples from Chile (12.6 %) and Germany (29.4 %). The high abundance of *Crenarchaeota* seen here in samples from Germany (Fig. S1) has also been identified by Kemnitz, Kolb and Conrad (2007), but in their case in samples of an acidic forest soil. In contrast to the present observation, studies of soil samples from Chilean vineyards have previously reported relative abundances of *Crenarchaeota* below 0.1 % (Castañeda and Barbosa 2017). For *Crenarchaeota*, the ammonia-oxidising archaea (AOA) *Nitrososphaera* was the main genus detected in all the countries surveyed in this study. This genus has been observed by Zhalnina et al. (2013) to significantly respond to agricultural management practices. In the present study this taxon was also one of the best predictors from the random forest model.

Of the fungi, *Solicoccozyma* (mostly known as *Cryptococcus*) was the dominant genus in eight out of the 13 countries and the best predictor in the random forest model considering fungal community alone. It belongs to the group of oxidative basidiomycetous yeasts and has been found to be associated with the phyllosphere, grapes and soil (Barata, Malfeito-Ferreira, and Loureiro 2012). *Cryptococcus*, *Saccharomyces* (Spain and USA) and *Candida*, *Hanseniaspora* and *Pichia* (these last three genera were not identified in the samples in the present study) provide most of the diversity of the frequently isolated yeast species related to grape or isolated from fermented grape juice (Kachalkin et al. 2015). Studies have shown that *Cryptococcus* are not severely affected by fungicides, which may explain their higher abundance in vineyards (Čadež, Zupan, and Raspor 2010; Comitini and Ciani 2008). Some of the other main fungal taxa identified in the present samples also contained documented plant pathogens at a lower resolution level. *Fusarium*, detected among the dominant taxa in Australia, Denmark, Germany Portugal and South Africa, is recognised as containing many plant pathogenic and fruit spoilage species (Kepler, Maul, and Rehner 2017). *Gibberella intricans*, which was found in three countries above 2 %, is a teleomorph (i.e. the sexual reproductive stage) of *Fusarium equiseti*, a cosmopolitan soil fungus associated with stem and root rot in different plant species (Pitt and Hocking 2009). Pancher et al. (2012) isolated the species *Gibberella baccata* (*Fusarium sambucinum*) from the grapevine endosphere of samples from Trentino, Northern Italy, while in the present study it appeared in samples from Denmark and Germany (also sharing a relatively cool climate with the Trentino region of Italy). *Cladosporiaceae*, and within this family *Cladosporium*, also widely detected in the present samples (especially in Denmark, where it reached 19 %), contain several species identified as fungal pathogens on grapevines (Jayawardena et al. 2018). However, the detection of these potential phytopathogens does not directly indicate plant infection because the success of microorganisms to affect plant health relies on whole soil and rhizosphere microbial interactions (Berendsen, Pieterse and Bakker 2012) and strain-specific virulence factors that cannot be retrieved with amplicon sequencing. Their existence in soil can be due to the fact they are present as spores or dead cells. Furthermore, it is not possible to infer pathogenicity based on a phylogenetic relation to a pathogen.

Based on the random forest results, both 16S and ITS abundances showed good predictive power for the geographical region. Overall, accuracies were 83 % and 86 % on a national scale for the prokaryotic and fungal communities respectively (Table S5). These results included the list of the best predictor taxa, some of which are also known to play an ecological role in vineyards, such as *Nitrososphaera* and *Cryptococcus*, which also appeared as the most dominant taxa within the prokaryotic and fungal core communities (Fig. 3b). The few occasions on which the confusion matrix gave a misprediction could be attributed to several factors; one was due to the approximation applied to overcome the different coverage of the US states represented in the dataset. In fact, although all the samples from the United States of America were considered to be from a single country, they were collected in three different states (California, Oregon and Texas) which increased the dispersion and group variance fooling the model. Finally, apart from all the regional signatures described above, it should be also highlighted that a core microbiome could be identified across vineyard soils, as previously described in other global surveys (Delgado-Baquerizo et al. 2018; Egidi et al. 2019)

## 5 Conclusions and perspectives

This study has provided new insights into microbial *terroir*, quantifying it on different scales for the first time. This concept was extended to a global scale, showing a hierarchical effect that is valuable on continental, national and regional scales. The level of resolution reached here, together with some other evidence reported at a local (inter-block and intra-block) scale (Alonso et al. 2019), suggests that the microbiome should be considered as an important variable in identifying agricultural sites for the definition of homogeneous functional zones such as basic *terroir* units. These should represent the smallest area for which it is possible to objectively describe the effect of environment on plant physiology and agricultural production, and which could be differentially managed. This is the first step towards achieving precision and a sustainable agriculture framework. Since the microbial *terroir* appears to be dependent on several different factors, from geography to climate, soil characteristics and vineyard management, there is a thread that links them all and this must be sought in the hidden dynamics of their microbial communities. Thus, the use of microbial information as a way of discriminating between vineyards in different countries provides the first applicative use of this technology and tools for improving the accuracy and representativeness of the microbial map by adding new samples in future. The random forest model developed ultimately confirmed this study’s initial hypothesis that geography determines the microbiota to such an extent that it can be used to predict the origin of a vineyard’s soil. This is therefore another argument supporting the definition of appellations of origins, both in legal and marketing terms. It also supports its possible application to other products with an historical attachment to the concept of *terroir* (e.g. cheese, coffee, cocoa etc.), using the microbiome to complement the evidence of other chromatographic and sensory analytical techniques already used to address wine typicality (Cadot et al. 2010; Maitre et al. 2010). Finally, these results should encourage further explorations of the significance and limits of the microbial aspect of agricultural *terroirs*, since they provide a baseline for guiding future studies in the field. Such studies should include observations of temporal and seasonal dynamics but should also follow a more functional approach (using metagenomics/metatranscriptomics to disentangle soil food webs) for a better understanding of the effect of microbial *terroir* on the health status of vineyard soils and how human intervention may exploit microbial dynamics in vineyards to improve the functional biodiversity of vineyard soils.

## Conflict of Interest

A.Acedo, and R. Ortiz-Álvarez are currently employed at Biome-Makers. I. Belda was employed by Biome Makers (contributing to this work under the framework of his postdoctoral Torres Quevedo Grant – PTQ08253) but currently working at Complutense University of Madrid (Spain).

## Acknowledgements

The authors would like to thank the Horizon 2020 Programme of the European Commission within the Marie Schlodowska-Curie Innovative Training Network “MicroWine” (grant number 643063) for financial support and all the wineries that provide samples to perform this study.

## Notes

### Competing Interest Statement

A.Acedo, and R. Ortiz Alvarez are currently employed at Biome Makers. I. Belda was employed by Biome Makers (contributing to this work under the framework of his postdoctoral Torres Quevedo Grant PTQ08253) but currently working at Complutense University of Madrid (Spain).

