## Supplementary Figures for "Microbial map of the world’s vineyards: Applying the concept of microbial *terroir* on a global scale"

* IB and LHH supervised the project equally

¥ corresponding author

**Supplementary Information**

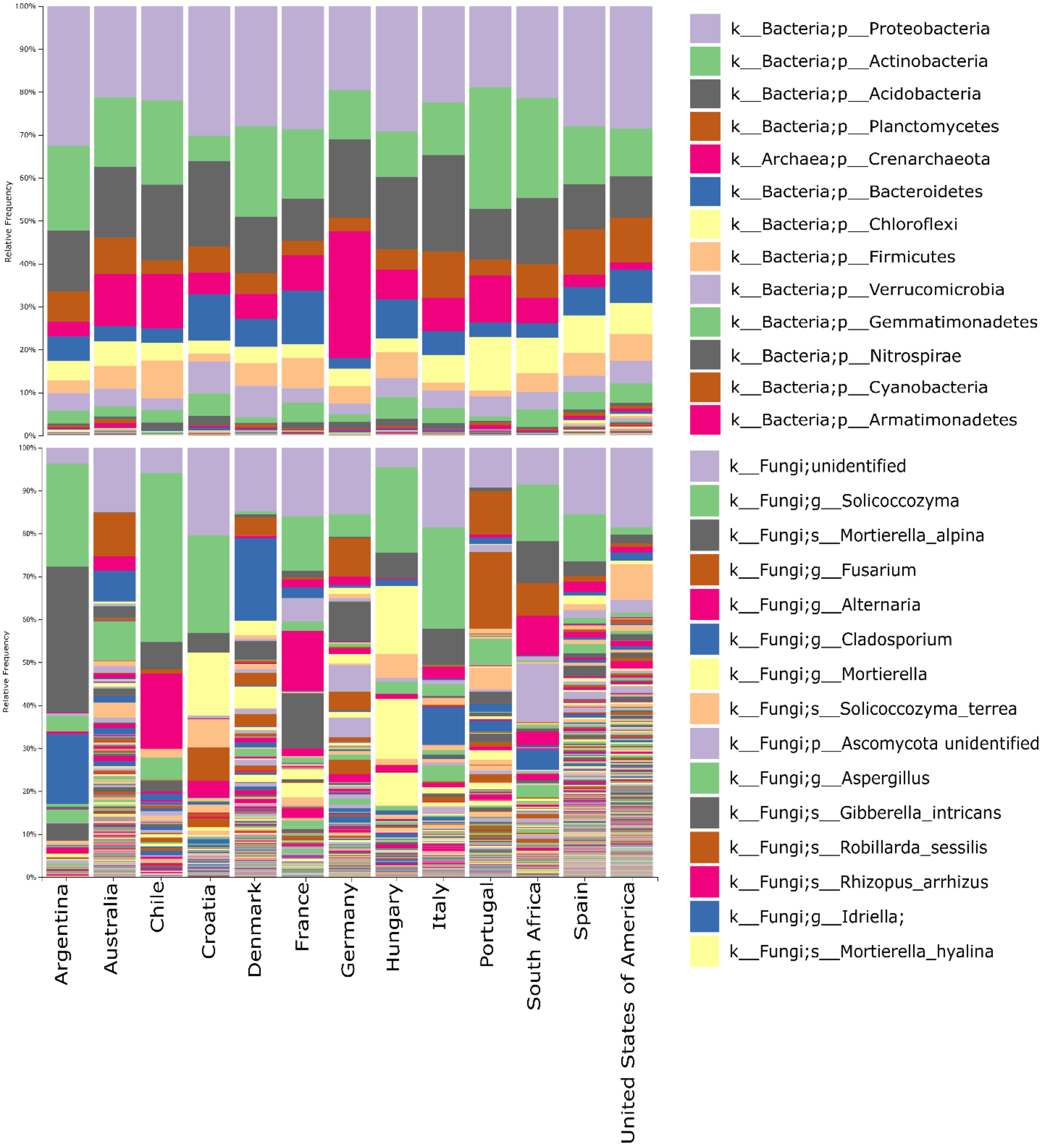


Figure S1. Taxonomical composition at country level: above) a phyla-level taxa bar plot representing the samples' 16S community, grouped by country; the colours identify the 13 most represented phyla. Below) a taxa bar plot representing the fungal community at the highest rank for the 15 most represented taxa in the samples, grouped by country.

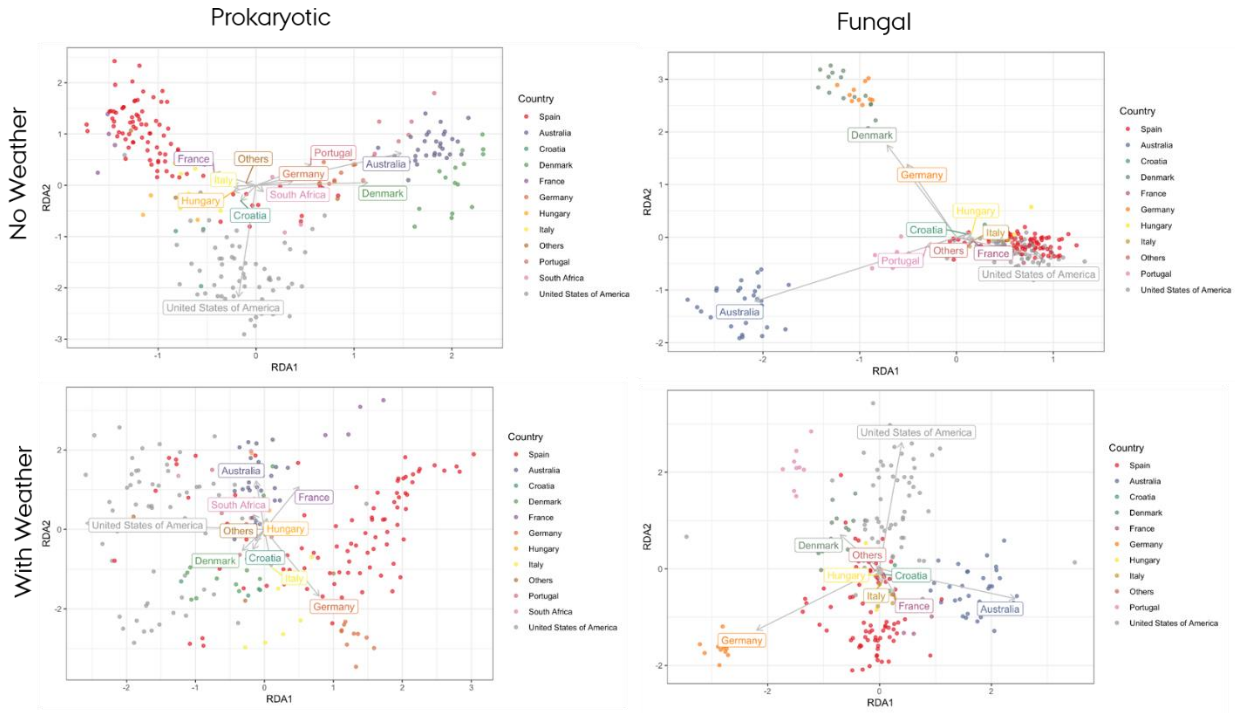


Figure S2. Variance partitioning RDA on the prokaryotic and fungal community with and without weather conditions. Samples are coloured based on the countries they come from.

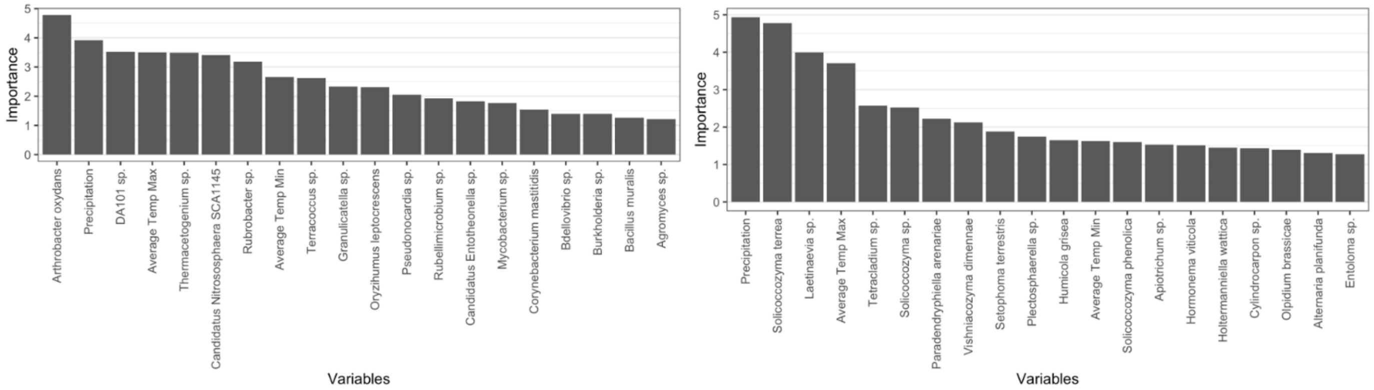


| | Australia | Denmark | Germany | Spain | United States of America | Others |
|---|---|---|---|---|---|---|
| *Australia* | 6 | 0 | 0 | 0 | 0 | 0 |
| *Denmark* | 0 | 4 | 0 | 0 | 0 | 0 |
| *Germany* | 0 | 0 | 1 | 0 | 0 | 0 |
| *Spain* | 0 | 0 | 0 | 19 | 0 | 2 |
| *United States of America* | 0 | 0 | 0 | 3 | 14 | 0 |
| *Others* | 0 | 0 | 0 | 7 | 0 | 4 |

| | Australia | Denmark | Germany | Spain | United States of America | Others |
|---|---|---|---|---|---|---|
| *Australia* | 6 | 0 | 0 | 0 | 0 | 0 |
| *Denmark* | 0 | 2 | 0 | 0 | 0 | 0 |
| *Germany* | 0 | 0 | 3 | 0 | 0 | 0 |
| *Spain* | 0 | 0 | 0 | 24 | 0 | 1 |
| *United States of America* | 0 | 0 | 0 | 2 | 12 | 1 |
| *Others* | 0 | 0 | 0 | 2 | 0 | 5 |

Figure S3. Figure 4. Random forest results at national scale including microbial and meteorological data. The top charts shows the confusion matrixes for the prokaryotic and fungal communities; the bottom charts give the 20 best predictor for prokaryotic (right) and fungal (left) community.
